# Notch Signaling in regulating Bone-derived Nanoparticles (BNPs) enhanced Osteogenic Differentiation

**DOI:** 10.1101/2024.06.19.599760

**Authors:** Austin Stellpflug, Justin Caron, Samantha Fasciano, Bo Wang, Shue Wang

**Author notes:** Corresponding Author: Dr. Bo Wang is with Marquette University and the Medical College of Wisconsin, Milwaukee, WI 53226, USA, Dr. Shue Wang is with the University of New Haven, West Haven, CT, 06516, USA.

## Abstract

Mesenchymal stem cells (MSCs)-based bone tissue regeneration has gained significant attention due to their excellent differentiation capacity and immunomodulatory activity. Enhancing osteogenesis regulation is crucial for improving the therapeutic efficacy of MSC-based regeneration. By utilizing the regenerative capacity of bone ECM and the functionality of nanoparticles, we recently engineered bone-based nanoparticles (BNPs) from decellularized porcine bone. The effects of internalization of BNPs on MSCs viability, proliferation, and osteogenic differentiation were first investigated and compared at different time points. The phenotypic behaviors, including cell number, proliferation, and differentiation were characterized and compared. By incorporating this LNA/DNA nanobiosensor and MSCs live cell imaging, we monitored and compared Notch ligand delta-like 4 (Dll4) expression dynamics in cytoplasm and nucleus during osteogenic differentiation. Pharmacological interventions are used to inhibit Notch signaling to examine the mechanisms involved. The results suggest Notch inhibition mediates osteogenic process, with reduced expression of early and late stage of differentiation markers (ALP, calcium mineralization). The internalization of BNPs led to an increase in Dll4 expression, exhibiting a time-dependent pattern that aligned with enhanced cell proliferation and differentiation. Our findings indicate that the observed changes in BNP-treated cells during osteogenic differentiation could be associated with the elevated levels of Dll4 mRNA expression. In summary, this study provides new insights into MSCs osteogenic differentiation and the molecular mechanisms through which BNPs stimulate this process. The results indicate that BNPs influence osteogenesis by modulating Notch ligand Dll4 expression, demonstrating a potential link between Notch signaling and the proteins present in BNPs.

## Introduction

Bone marrow derived mesenchymal stem cells (MSCs) play a key role in bone repairing and regeneration by differentiation to bone-forming osteoblasts and cartilage-forming chondrocytes ^1–5^. MSCs contribute to bone healing through three different approaches: a) differentiation and replacement; b) secretion of cytokines and extracellular vesicles; and c) immunomodulatory activity ^6–8^. While identifying the most efficient method for enhancing osteogenic differentiation of MSCs to promote bone regeneration remains challenging, regulating osteogenesis is crucial for improving the therapeutic efficacy. While the differentiation potential of MSCs is well-established, the specific mechanisms governing their plasticity are not fully understood, particularly the processes by which MSCs differentiate into osteoblasts for bone formation. Over the last few decades, unremitting efforts have been devoted to understanding biochemical signals that regulate MSCs commitment. Based on these efforts, a number of chemical stimuli (e.g., small bioactive molecules, growth factors, and genetic regulators) have been identified in regulating MSCs lineage commitment, including bone morphogenetic protein (BMP), Wnt, and Notch signaling ^9–11^. Enhancing osteogenesis is crucial for improving the effectiveness of MSC-based therapies in bone tissue engineering and regeneration. Thus, recent research has focused on developing strategies to enhance osteogenesis, which involve biophysical and biochemical stimulation. In recent years, the rapid advancements in nanotechnology and nanomedicine have significantly transformed regenerative medicine, especially in the context of bone diseases ^12–14^. Nanoparticles (NPs) have emerged as multifunctional tools that integrate diagnostic and therapeutic functions, offering new avenues for treatment ^15–19^. Moreover, NPs offer precise control over stem cell behavior and enhance drug delivery by overcoming biological barriers due to their small size. They also allow for prolonged circulation and retention of therapeutic agents, thereby reducing the side effects associated with traditional methods.

NPs can be broadly classified into inorganic particles (such as ceramics, metal, silica, gold, and silver) and organic particles (including synthetic polymers, liposomes, and proteins) based on their chemical structures ^20^. Although the therapeutic applications of NPs in clinical settings have been extensively studied, several challenges remain in translating these findings into practical treatments. Key issues include their low biocompatibility and the risk of inducing inflammation and tissue damage, which have hindered the wider adoption of NPs in clinical practice, highlighting the need for the development of safer and more effective alternatives ^21–23^.

Our lab has recently developed a novel type of bone-derived nanoparticles (BNPs) from decellularized porcine bones ^24^. These BNPs have shown promising bone regenerative potential both *in vitro*, by promoting osteogenic differentiation; and *in vivo*, by repairing bone defects when used as graft material. The utilization of BNPs offers a novel approach to overcoming the limitations of traditional NPs, which will provide a more biocompatible and efficient delivery system for therapeutic agents.

Despite these findings, the fundamental mechanisms through which BNPs affect osteogenic differentiation process remain unexplored. Understanding these mechanisms is crucial for optimizing the design and application of BNPs in bone regeneration. Osteogenic differentiation is a dynamic process and involves several significant signaling pathways, including BMP signaling, Wnt/b-catenin signaling, Hedgehog (HH) signaling, and YAP/TAZ (transcriptional coactivator with PDZ-binding motif), and Notch signaling ^25, 26^. Our recent studies have shown Notch signaling is required and modulate shear-stress induced osteogenic differentiation ^2^. In addition, current studies revealed internalization of NPs could enhance osteogenic differentiation ^27–29^. However, the involvement of Notch signaling remains obscure due to a lack of effective tools to detect and monitor the gene expression in live cells. Traditional methods for gene detection are limited due to the need of isolation or fixation, which results in the loss of spatial and temporal gene information. Techniques such as RNA in situ hybridization and single cell transcriptomics only apply to fixed cells, thus limiting their utility ^30^. Although fluorescent protein tagging allows for the monitoring of RNA dynamics in live cells, it is restricted by low transfection efficiency and the requirement for genetic modifications to express engineered transcripts ^31^. Thus, dynamic monitoring of gene expression in live cells at the single cell level will reveal the fundamental regulatory mechanism of cells during dynamic biological processes, which will eventually open opportunities to develop novel approaches for tissue engineering and regenerative medicine.

Here, we utilized a double-stranded locked nucleic acid/DNA (LNA/DNA) nanobiosensor to investigate the regulatory role of Notch signaling during BNPs induced osteogenic differentiation. The effects of internalization of BNPs on MSCs viability, proliferation, and osteogenic differentiation were first investigated and compared at different time points. The phenotypic behaviors, including cell number, proliferation, and differentiation were characterized and compared. By incorporating this LNA/DNA nanobiosensor and live cell imaging, we monitored and compared Notch ligand delta-like 4 (Dll4) expression dynamics in cytoplasm and nucleus during osteogenic differentiation. Pharmacological interventions are used to inhibit Notch signaling to examine the molecular mechanisms involved. The results suggest Notch inhibition mediates osteogenic process, with reduced expression of early and late stage of differentiation markers (ALP, calcium mineralization). The internalization of BNPs led to an increase in Dll4 expression, exhibiting a time-dependent pattern that aligned with enhanced cell proliferation and differentiation. Our findings indicate that the observed changes in BNP-treated cells could be associated with the elevated levels of Dll4 mRNA expression. In summary, this study provides new insights into MSCs osteogenic differentiation and the molecular mechanisms through which BNPs stimulate this process. The results indicate that BNPs promote osteogenesis by modulating Notch ligand Dll4 expression, demonstrating a link between Notch signaling and the proteins present in BNPs. Future research will explore the interactions between TGF-β, Notch, and BMP signaling pathways and the impact of BNPs on these interactions during osteogenic differentiation.

## Materials and Methods

### Fabrication of bone derived nanoparticles

The BNPs were fabricated following our previously published protocol ^24^. Briefly, fresh porcine tibias were fully demineralized with 0.5 M hydrochloric acid (HCL, Sigma-Aldrich) and decellularized with 0.5% sodium dodecyl sulfate (SDS) and 1% Triton X-100 mixed solution (Sigma-Aldrich). To prepare the digested bone solution, the demineralized and decellularized bones were freeze-dried at −80 °C, milled into a fine powder, and digested in HCL solution (1 gram of extracellular matrix (ECM) powder/100 ml of 1M HCL) 15% w/w pepsin under constant stirring at 45 °C until the whole solution became a smooth, uniform liquid without visible solid particles, indicating that the digestion process was complete ^24^.

To fabricate the BNPs, the protein concentration of the final digested bone ECM solution was adjusted to a final concentration of 330 µg/mL prior to desolvation. Acetone was added dropwise to the ECM solution (the volume ratio (mL) of acetone to ECM solution was 3:1) under constant stirring at room temperature. 10 minutes after the last of the acetone was added to the ECM solution, 50% glutaraldehyde solution was added in a ratio of 33 µL of glutaraldehyde per 1 mL of the initial ECM solution only. This solution was left to stir at room temperature for 30 minutes, upon which the pellet of BNPs was collected via centrifugation. The BNPs were then washed with distilled water three times, resuspended again into distilled water, and dispersed via sonication. The BNPs were then freeze-dried at −80 °C and used for experimental characterization or long-term storage.

### Characterization of BNPs

Morphology of the BNPs was characterized using scanning electron microscopy (SEM). BNP samples were resuspended into DI water, sonicated, and 100µL aliquots placed on copper mesh grids, being left to dry for 24 hours to remove excess moisture. Samples were sputter coated with gold-palladium and observed with a JEOL JSM 35 scanning electron microscope (JEOL USA, Inc., Peabody, MA).

### Cell culture and reagents

Bone marrow derived MSCs (Lonza) were cultured in MSCBM (Catalog #: PT-3238, Lonza) with GA-1000, L-glutamine, and mesenchymal cell growth factors (Catalog #: PT-4105, Lonza). Cells were cultured and maintained in a tissue culture dish at 37 ℃ and 5% CO_2_ in a humidified incubator. The medium was replaced every two days and cells were passaged using 0.25% Trypsin-EDTA (Invitrogen). Cells were seeded at 0.1×10 ^6^ and 0.05×10 ^6^ cells/well for 12 and 24 well plates, respectively. The newly seeded cells were cultured until reach 80% confluency before induction medium was added. Cells from passage 2-6 were used in the experiments. For intracellular uptake of BNPs, BNPs were re-suspend in opti-MEM (ThermoFisher) and sonicated for 1 minute using probe sonicator (Brandon). Unless specified, cells were incubated with BNPs at a concentration of 20 µg/mL overnight. Osteogenic induction medium (Catalog #: PT-3002, Lonza) were added and replaced every three days. For control groups, cells were maintained in basal medium without induction. To investigate the involvement of Notch signaling, cells were administrated with γ-secretase inhibitor DAPT (20 µM) on cell differentiation. It is noted that cells without DAPT treatment were added DMSO for control. The differentiation effects were accessed and compared after 7, 14, and 21 days of induction.

### Design and preparation of LNA probe

A LNA/DNA nanobiosensor is a complex of a detecting probe and a quencher ^32^. The detecting probe is a 20-base nucleotide with alternating LNA/DNA monomers, which sequence is complementary to target mRNA sequence. A fluorophore (6-FAM) was labeled at the 5’ of detection probe for visualization of mRNA signal in cells. The probe design process has been reported previously, which includes acquiring target mRNA sequence from GeneBank, choosing a 20-base pair ^33–35^. The selected nucleotide sequence will be characterized and optimized for stability and specificity using mFold server and NCBI Basic Local Alignment Search Tool (BLAST) database, respectively. The quencher probe consists of a 10-base pair nucleotide sequence, incorporating LNA/DNA monomers, designed to be complementary to the 5’ end of the LNA detection probe. At the 3’ end of this quencher probe, an Iowa Black RQ fluorophore is attached for labeling purposes. The Dll4 LNA detecting probe was designed based on target mRNA sequences (5’-3’: +AA +GG +GC +AG +TT +GG +AG +AG +GG +TT). All the probes were synthesized by Integrated DNA Technologies Inc. (IDT).

To assemble LNA/DNA nanobiosensor, the LNA detecting probe and quencher probe were first dissolved in 1x Tris EDTA buffer (pH=8.0) at a concentration of 100 nM. These components were combined in a 1:2 ratio and heated to 95 ℃ for 5 minutes in a dry water bath, then allowed to cool to room temperature gradually over 2 hours. Once cooled, the mixture can be refrigerated and stored for up to 7 days. For mRNA detection, this nanobiosensor was transfected into cells using Lipofectamine 2000 (ThermoFisher) according to the manufacturer’s instructions.

### Cell proliferation and viability

The cell proliferation and viability after BNPs incubation was evaluated using cell counting kit-8 (cck-8, Sigma Aldrich), following the manufacturers’ instructions. Cells were seeded at a concentration of 200 cells/well in 96-tissue culture well plates with the volume of 100 µL culture medium. After 24 hr incubation, cells were treated with BNPs at different concentrations (10, 20, 50, 100 µg/mL) after 14 days of incubation. The proliferation was evaluated at day 1, 2, 3, 5, and 10, respectively. After incubation, cells were added with cck-8 and cultured for 4 hr. The absorbance was measured at 450 nm for each sample and compared using a fluorescence microplate reader (BioTek, Synergy 2).

### Live/Dead viability staining

The cell viability was further evaluated using live/dead viability assay (ThermoFisher). Cells were stained using propidium iodide (PI, 10 μg/mL), a fluorescent agent that binds to DNA by intercalating between the bases with little or no sequence preference. Hoechst 33342 was used to stain cell nucleus at a concentration of 20 μM for 30 minutes. After staining, cells were washed three times with 1x PBS to remove extra dye. Cells were then imaged using Texas Red (535/617 nm) and DAPI (360/460 nm) filters using Echo Revolution Fluorescent Microscope.

### ALP and Alizarin Red S Staining

To evaluate osteogenic differentiation, the expression of early differentiation marker (alkaline phosphatase, ALP) and late-stage differentiation marker (calcium mineralization) were quantified using ALP staining kit (Sigma Aldrich) and Alizarin Red S (ARS) staining quantification assay (ScienCell), respectively. For ALP staining, the staining solution was prepared by combining Fast Red Violet solution, Naphthol AS-BI phosphate solution, and water in a 2:1:1 ratio. Subsequently, cells were fixed with 4% cold Paraformaldehyde (PFA) for 2 minutes to preserve ALP activity. After fixation, the PFA was removed without washing the cells. The staining solution was then added to the fixed cells for 15 minutes at room temperature and protected from light. Following staining, the cells were washed three times with 1x PBS before imaging.

For ARS staining, the culture medium was removed, and the cells were washed with 1xPBS for 3 times. The cells were then fixed with 4% PFA for 15 minutes at room temperature. Then the PFA was removed, and the cells were washed 3 times with diH_2_O. 40 mM ARS solution was added to each well. The wells were shaken gently for 20-30 minutes with the stain. Lastly, the stain solution was removed, and the cells were washed five times with diH_2_O. Images were then taken of the stained cells.

### Imaging and statistical analysis

Images were obtained using Echo Revolution fluorescent microscope with an integrated digital camera (5MP CMOS Color for bright field, 5 MP sCMOS Mono for fluorescence imaging). To ensure consistency, all images were captured under identical settings, including exposure time and gain. Image analysis and data collection were conducted using NIH ImageJ software. To measure Dll4 mRNA and ALP enzyme activity, the mean fluorescence intensity of each cell was measured, and background noise subtracted. Cells were quantified within the same field of view, with a minimum of five images analyzed for each condition. The experiments were conducted at least three times, with over 100 cells quantified per group. Results were analyzed using independent, two-tailed Student t-test in Excel (Microsoft). P < 0.05 was considered statistically significant.

## Results

### Fabrication of BNPs and mechanisms of double-stranded LNA nanobiosensor

The BNPs fabricated were consistent in morphology with our previous findings ^24^. **Fig. 1A** showed the fabrication process, including bone decellularization and demineralization, lyophilized bone milling, digesting and desalting bone ECM powder, and final synthesis. The synthesized BNPs has an average size of 79 ± 25.4nm, **Fig. 1B**. A double-stranded LNA nanobiosensor was utilized to investigate the involvement of Notch signaling during the dynamic osteogenic differentiation process. The double-stranded DNA/LNA nanobiosensor is a complex of an LNA detecting probe and an LNA quenching probe, with the length of 20 and 10 nucleotides, respectively, **Fig. S1**. The LNA detection probe is a single-stranded oligonucleotide sequence with alternating LNA/DNA monomers, which are designed to be complementary to the target mRNA sequence. The LNA nucleotides were chosen due to their higher thermal stability compared to DNA nucleotides, thus enhancing the specificity and sensitivity. To visualize the mRNA expression in live cells, a fluorophore (6-FAM (fluorescein)) was labeled at the 5’ end of the detecting probe. The design process of this nanobiosensor have been reported previously. Briefly, the detecting LNA probe will spontaneously binds to quenching probe, forming an LNA-quencher complex. This proximity allows the quencher to quench the fluorescence of the fluorophore at the 5’ end of the LNA probe due to quenching properties. The LNA-quencher complex will be transfected into cells for mRNA detection and visualization. Upon cellular uptake, the LNA detecting probe disassociates from the quencher and binds to target mRNA molecules, thus reacquiring a fluorescence signal. This displacement is due to a greater binding free energy difference between the LNA probe and target mRNA compared to the LNA probe and quencher. Consequently, the fluorescence intensity within individual cells that contain the LNA/DNA nanobiosensor can quantitatively measure the amount of target mRNA present. In this study, MSCs were transfected with the LNA/DNA nanobiosensor before osteogenic induction.

**Fig. 1.**
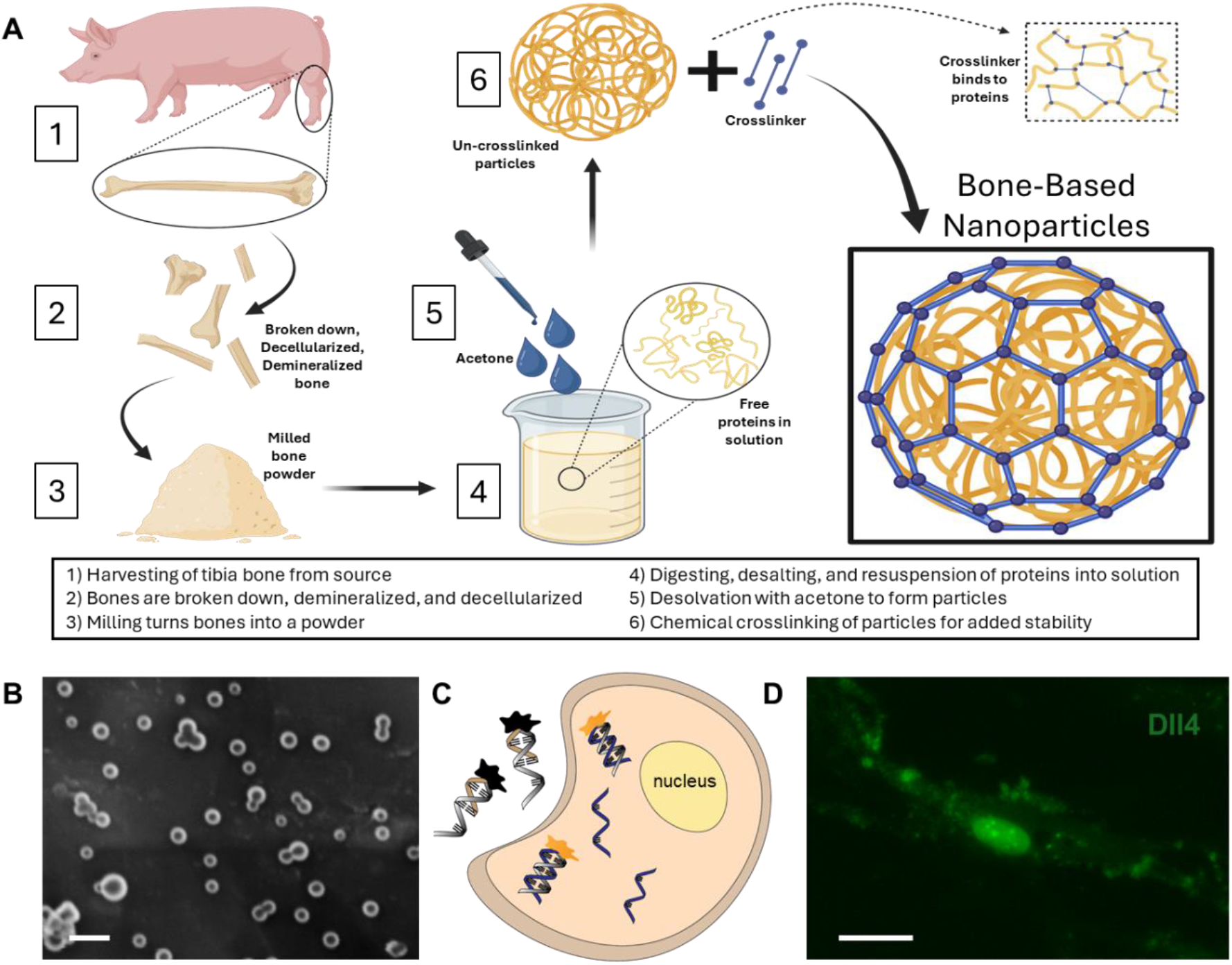
BNPs fabrication process and locked nucleic acid (LNA)/DNA nanobiosensor for investigation of Notch signaling in osteogenic differentiation. **(A)** Illustration of the fabrication process, including bone decellularization and demineralization, lyophilized bone milling, digesting and desalting bone ECM powder, and final synthesis. **(B)** SEM images of BNPs. Scale bar: 100 nm. **(C)** Biosensing mechanism of the LNA/DNA nanobiosensors. The detecting LNA probe, a 20-base nucleic acid molecule labeled with a fluorophore (6-FAM (fluorescein)) at the 5′ end, binds to quencher probe to quench fluorescence signal. After internalization by MSCs, LNA probes bind targeted mRNA in the cytoplasm and reacquire fluorescence. The fluorescence intensity thus serves as an indicator of the expression level of target mRNA. **(D)** Dll4 mRNA expression in single MSC with BNPs treatment (20 µg/mL). Scale bar: 10 µm.

### BNPs enhance MSCs proliferation

In order to study the effects of different concentrations of BNPs on cell proliferation and viability, MSCs were cultured with BNPs at the concentration of 0, 5, 10. 20, and 50 µg/mL for 24 hrs. The cell viability was first evaluated and compared using cck-8 kit (Sigma) after 14 days of incubation. **Fig. S2** shows the comparison of absorbance with and without BNPs incubation at different concentrations. Moreover, we evaluated the cell survival using live/dead cell assay after 5 and 7 days, respectively. **Fig. 2A** shows the bright field and fluorescent images of MSCs after 5 days of culture under control and BNPs co-culture group, respectively. It was evident that there are no dead cells in both control and BNPs co-culture group (third column of **Fig. 2A**), indicating BNPs co-culturing with MSCs did not affect the viability and cell survival. To further examine the effects of BNPs on cell proliferation, we quantified and compared the number of cells within different groups. **Fig. 2B** showed the comparison of quantified cell numbers with and without BNPs co-culturing after 5 and 7 days, respectively. The results showed that cell number increased 82.89 % and 57.36 % after 5 and 7 days, respectively. Since BNPs were fabricated using the entire ECM of decellularized porcine bone, BNPs contain a variety of ECM proteins ^24^. This data suggests that the effects of BNPs on cell proliferation occur at an early stage, indicating the ECM proteins were released quickly after several days of incubation. We further confirmed the effects of BNPs on cell proliferation after 1, 2, 3, 5, and 10 days of co-culturing. **Fig. 2C** shows the cumulative absorbance using cck-8 proliferation assay.

**Fig. 2.**
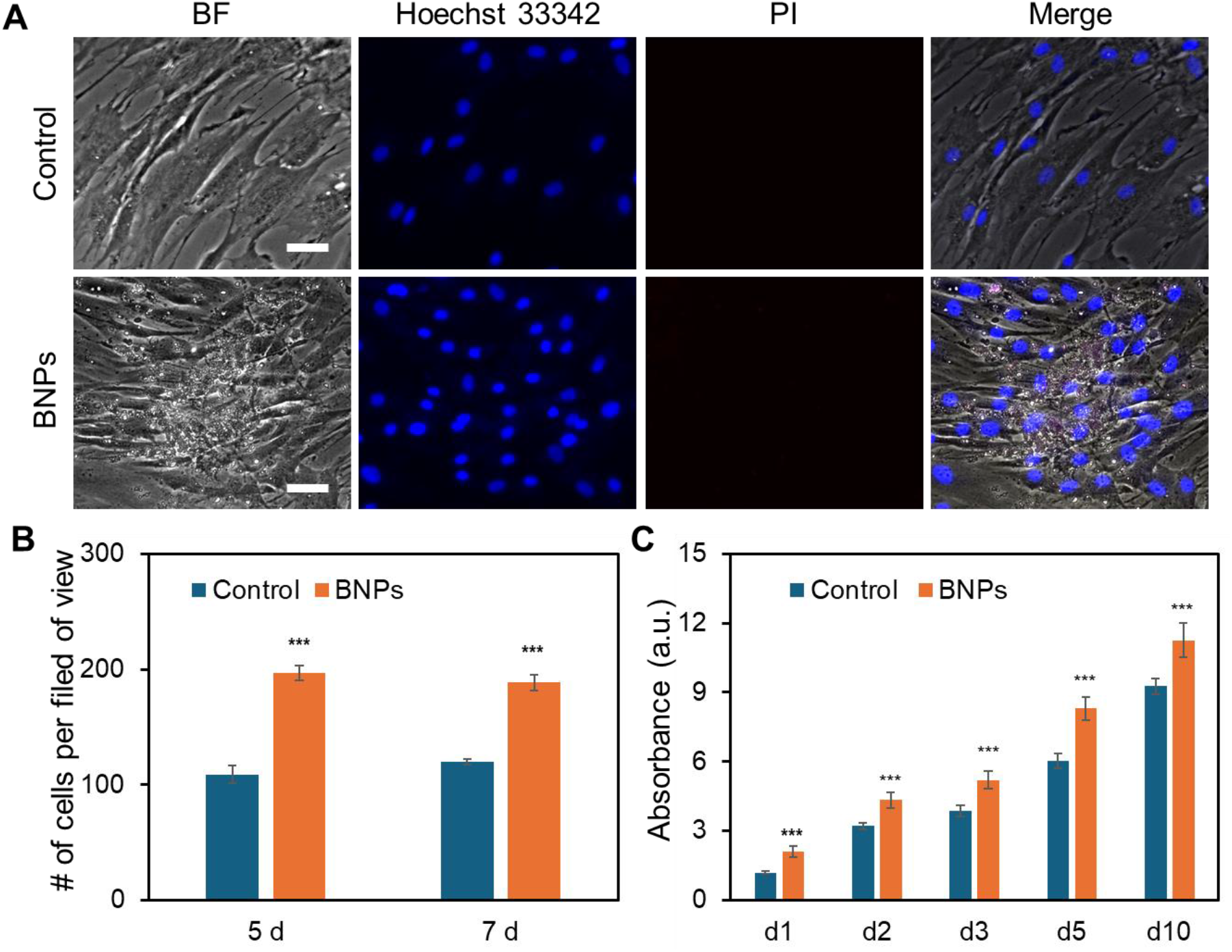
Effects of BNPs on MSCs viability. **(A)** Representative bright field and fluorescence images of MSCs under control and BNPs treated group after 5 days of incubation. Cells were stained with propidium iodide (PI, red), and Hoechst 33342 (blue), respectively. Scale bar: 50 μm. **(B)** Quantification and comparison of cell numbers with and without BNPs co-culturing after 5 days and 7 days, respectively. Cell numbers were calculated using a MATLAB program by counting the number of nucleuses in each field of view. At least 10 images were quantified for each condition. **(C)** Cumulative absorbance using cck-8 proliferation assay after 1, 2, 3, 5, and 10 days of incubation. Data are expressed as mean± s.e.m. (n=3, ***, P<0.001).

### Internalization of BNPs enhance osteogenic differentiation

We next evaluated the effects of BNPs on osteogenic differentiation by co-culturing BNPs with MSCs at the concentration of 20 µg/mL for 24 hrs. Briefly, MSCs were seeded in 12-well plates and cultured in the basal medium. Three groups of experiments were conducted, control (CTR group), osteogenic induction (OST group), BNPs and osteogenic induction (BNPs + OST group). Once the cells reach 70-80% confluency, BNPs were prepared in opti-MEM and added the cells in the BNPs + OST group, and osteogenic induction medium was replaced after 24 hrs of incubation. After 7 and 14 days of induction, osteogenic differentiation was evaluated using Alizarin Red S (ARS) Staining Quantification Assay (ScienCell). We further assessed calcium mineralization by extracting calcified minerals at low pH and neutralizing them with ammonium hydroxide. The calcium deposition was then quantified using colorimetric detection at 405 nm. **Fig. 3A** shows the bright field images of MSCs under different treatment after 7 and 14 days of incubation. **Fig. 3B** shows the images of different wells after ARS staining. The results showed that without osteogenic induction, there is no calcium deposition. With osteogenic induction but without BNPs incubation, calcium deposition was not visible after 7 and 14 days of induction, **Fig. 3A-3B**. It is evident that intracellular uptake of BNPs significantly enhanced osteogenic differentiation after 7 and 14 days, with visible calcium deposition. In order to investigate how BNPs accelerate osteogenic differentiation, we evaluated and compared ALP enzyme activity after 7 days of osteogenic induction. **Fig. 3C** showed representative fluorescence images of MSCs with and without BNPs internalization. We further quantified ALP activity by measuring the mean fluorescence intensity of ALP, **Fig. S3**. With BNPs internalization, the ALP activity increased by 1.5 folds compared to MSCs without BNPs. **Fig. 3D** shows the quantification and comparison results of calcium mineralization after 7 and 14 days of osteogenic induction, respectively. Without BNPs treatments, no calcium mineralization was observed for control, osteogenic groups after 7 and 14 days of induction. BNPs internalization significantly enhanced calcium mineralization, with approximately 5% and 23% of cells exhibit calcium mineralization after 7 and 14 days of induction, respectively.

**Fig. 3.**
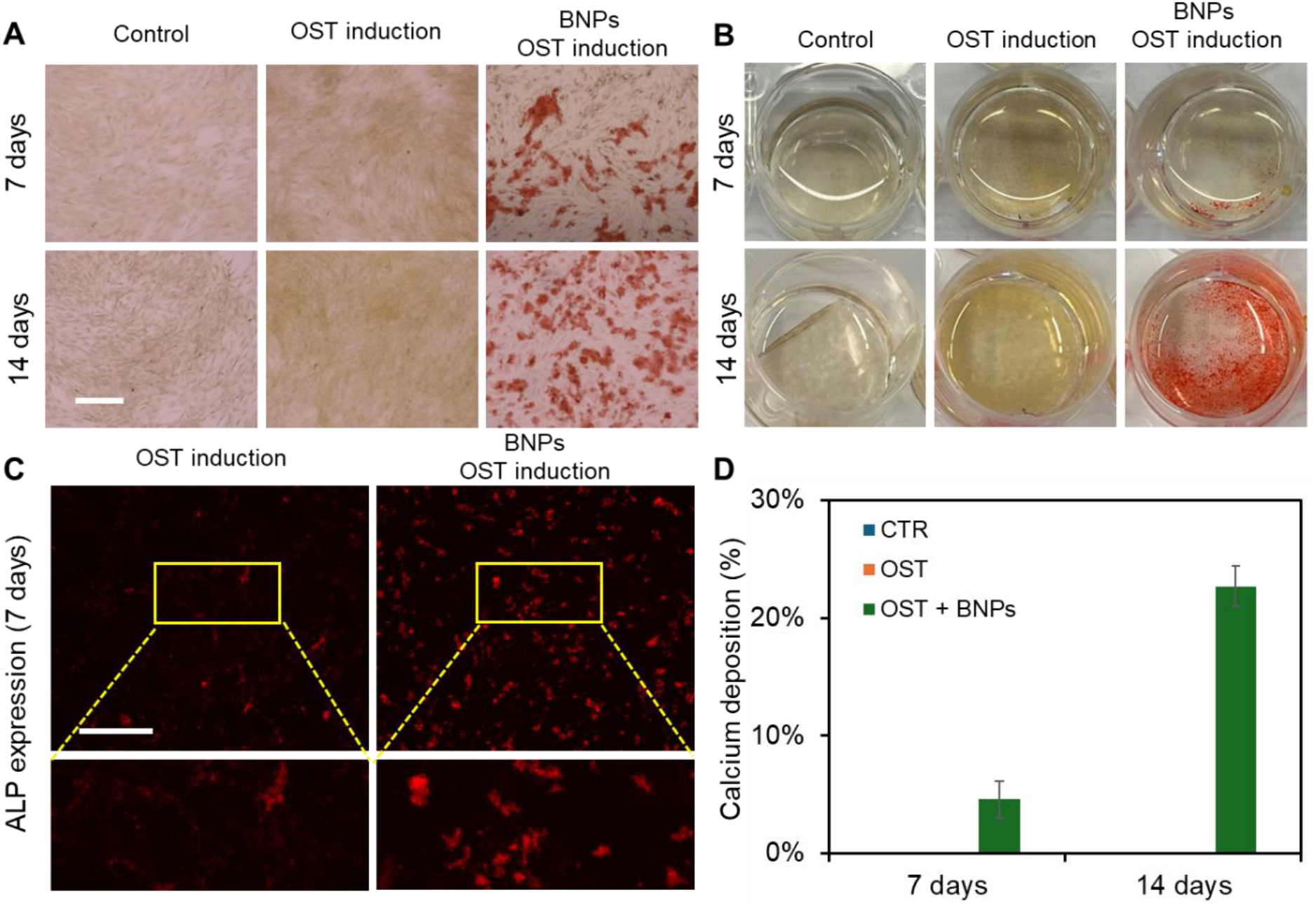
BNPs promote osteogenic differentiation of MSCs. **(A-B)** Representative images of MSCs under different treatments after 7 and 14 days of induction. CTR: control, cells were cultured in basal medium; OST: cells were induced using osteogenic induction medium; OST+BNP: Cells were treated with BNPs (20 µg/mL) and cultured in osteogenic induction medium. Scale bar: 200 µm. **(C)** Representative images of MSCs after 7 days of osteogenic induction under different groups. Scale bar: 200 µm. **(D)** Quantification of calcium mineralization after 7 and 14 days of osteogenic induction, respectively. The percentage of calcium deposition was quantified by measuring the ratio of ARS-stained cells to the total number of cells per field of view. A total of 10 images were quantified for each condition. expressed as mean± s.e.m. (n=5).

### Notch signaling is involved in BNPs induced osteogenic differentiation

Previous research has demonstrated that Notch signaling plays a role in the osteogenic differentiation, influencing ALP activity and the efficiency of osteogenic differentiation ^36–38^. Additionally, our group has recently identified Dll4 mRNA as a molecular biomarker of osteogenically differentiated MSCs ^4^. Inhibiting Notch signaling has been found to diminish osteogenic differentiation, accompanied by reduced ALP enzyme activity. However, it is obscure whether Notch signaling is involved in BNPs enhanced osteogenic differentiation. To better understand the regulatory role of Notch signaling, we utilized a pharmacological drug, γ-secretase inhibitor (DAPT) that blocks Notch endoproteolysis, to perturb Notch signaling. MSCs were treated with DAPT (20 µM) before osteogenic induction with or without BNPs incubation. The osteogenic differentiation across different treatments were evaluated and compared by quantifying ARS intensity, indicating the amount of calcium deposition. **Fig. 4A** and **Fig. S4** showed representative ARS staining images of MSCs after 21 days of osteogenic induction under different treatments. We next quantified the percentage of calcium deposition by measuring the average of ARS-stained region. **Fig. 4B** shows the comparison results of calcium deposition after 21 days of osteogenic induction. Without BNPs incubation, only about 5% of the cells exhibited calcium deposition, and DAPT treatment disturbed osteogenic differentiation, reducing calcium deposition to approximately 2%. Consistent with our earlier finding, treatment with BNPs significantly enhanced osteogenic differentiation, with approximately 34% of cells showing calcium deposition-an increase of about 4.6 times. Additionally, DAPT treatment reduced the effects of BNPs, with about 24% of cells exhibiting calcium deposition. Interestingly, DAPT treatment for the MSCs with BNPs has fewer effects on osteogenic differentiation, suggesting that the intracellular uptake BNPs counteracted the inhibitory effects of Notch signaling due to pharmaceutical treatment.

**Fig. 4.**
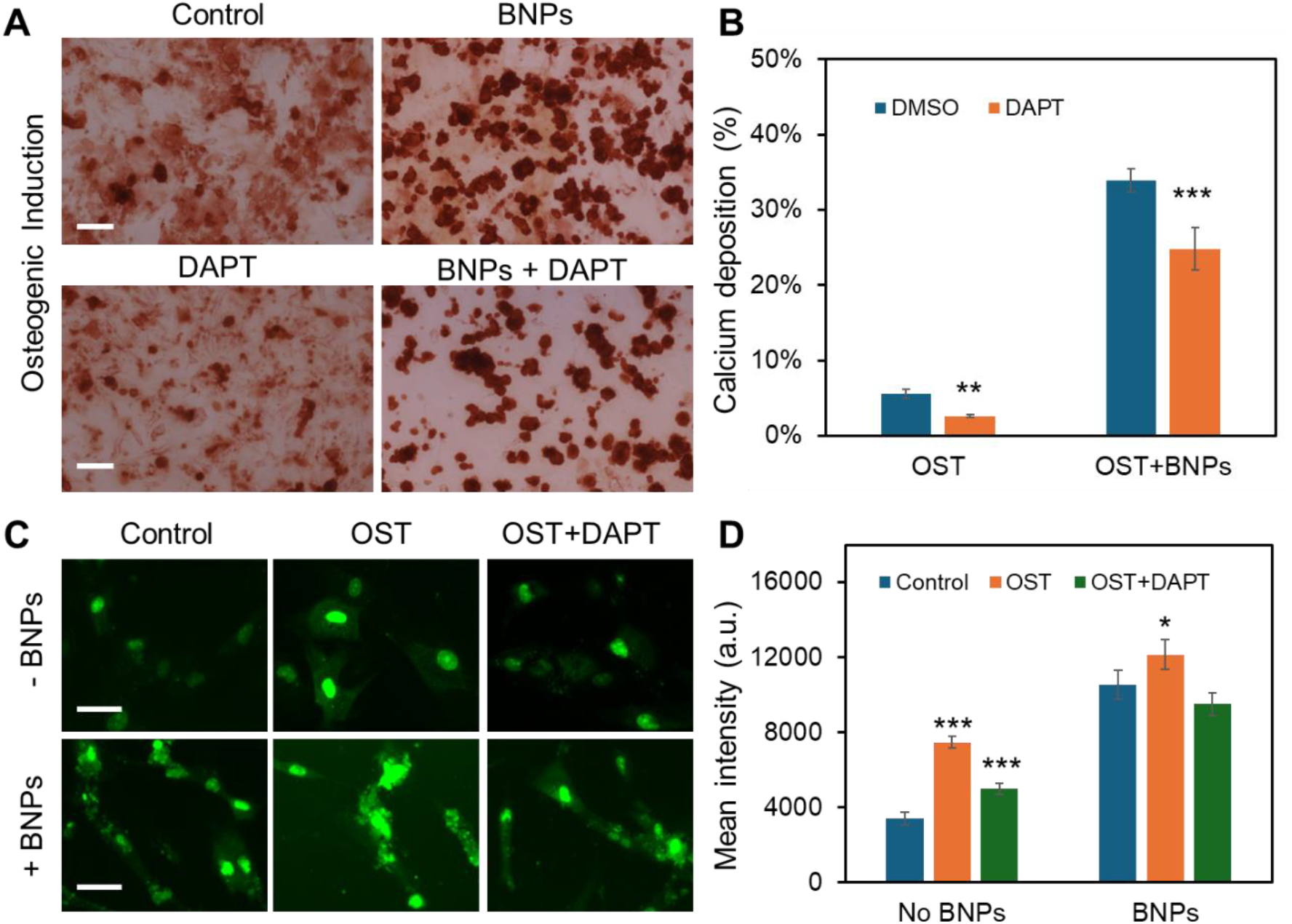
Notch signaling modulates osteogenic differentiation in both control and BNPs treated MSCs. **(A)** Representative bright field images of MSCs after days of osteogenic induction. For BNPs treated groups, MSCs were treated with BNPs at the concentration of 20 µg/mL overnight for internalization. DAPT (20 μM) were added to MSCs for Notch inhibition. Scale bar: 100 µm. **(B)** Quantification and comparison of calcium mineralization after 21 days of osteogenic induction. The percentage of calcium deposition was quantified by measuring the ratio of ARS-stained cells to the total number of cells per field of view. A total of 10 images were quantified for each condition. **(C)** Representative fluorescent images of MSCs under different treatment after 3 days of osteogenic differentiation. For the control group, cells were maintained in basal culture medium for comparison. DAPT was administered at a concentration of 20 µM. Green fluorescence signal indicates Dll4 mRNA expression. Scale bar: 50 µm. **(D)** Comparison of mean fluorescence intensity of Dll4 mRNA expression of MSCs after 3 days of osteogenic induction under different treatment. Data represent over 100 cells in each group and are expressed as mean± s.e.m. (n=3, ***, P<0.001, **, P<0.01, *, P<0.05)

To further explore the involvement of Notch signaling in the enhancement of osteogenic differentiation by intracellularly uptake of BNPs, we examined Notch 1 ligand, Dll4 mRNA expression of MSCs with or without BNPs incubation cultured in basal medium, induction medium, and induction medium with DAPT using an LNA/DNA nanobiosensor. **Fig. 4C** showed representative fluorescence images of MSCs with and without BNPs treatment under different conditions. Dll4 mRNA expression were quantified and compared by measuring the mean fluorescent intensity of each individual cells, **Fig. 4D**. The results showed that intracellular uptake BNPs enhanced Dll4 expression in all three groups, control, osteogenic, and DAPT treated osteogenic induction groups.

We also observed an increase in Dll4 expression following osteogenic induction, with a 1.2-fold increase in the control group and a 0.15-fold increase in the BNPs-treated group. In the BNPs-treated group, DAPT treatment moderated the enhancement of differentiation by BNPs, resulting in Dll4 mRNA expression that showed no significant difference compared to MSCs cultured in basal medium. These results suggest that Notch signaling plays a regulatory role in the osteogenic differentiation of MSCs in BNPs incubation. BNPs incubation increases Dll4 mRNA expression in MSCs undergoing osteogenic induction, highlighting Notch signaling’s role in BNPs regulated osteogenic differentiation. Furthermore, inhibiting Notch signaling diminishes the osteogenic differentiation induced by BNPs, resulting in decreased ALP enzyme activity and reduced Dll4 mRNA expression.

### Dynamic monitoring of Dll4 expression during osteogenic differentiation

To better understand and interpret the Notch regulatory mechanisms that contributed to osteogenic differentiation, we monitored Dll4 expression dynamics after osteogenic induction. The effects of BNPs incubation were assessed and compared with control (No BNPs) group. The capability of this LNA/DNA nanobiosensor of live-cell gene detection allows us to further elucidate the cytoplasmic and nucleus mRNA expression profile. **Fig. 5A** showed fluorescence images of MSCs expressing Dll4 after 1, 2, 3, and 5 days of osteogenic induction. To further explore and compare the distribution of Dll4 in cytoplasmic and nucleus, we quantified and compared Dll4 expression at different times with different treatments. **Fig. 5B** showed the comparison of cytoplasmic Dll4 expression with and without BNPs treatment. For the control group, the cytoplasmic Dll4 expression is relatively low compared to the BNP-treated group.

**Fig. 5.**
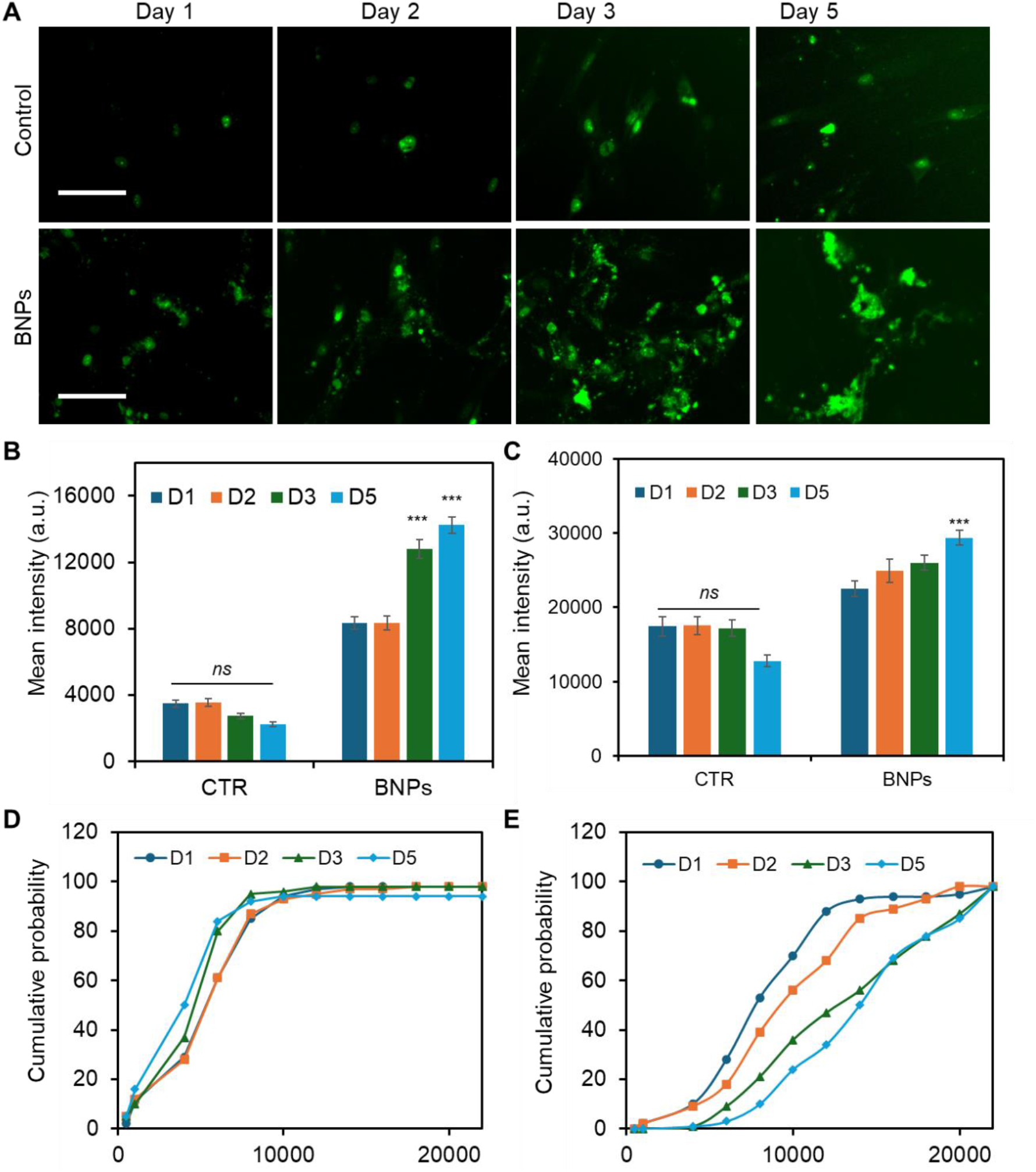
Dynamic single cell Dll4 mRNA expression analysis during osteogenic differentiation. **(A)** Representative fluorescence images of MSCs after 1, 2, 3, and 5 days of osteogenic induction. For BNPs group, MSCs were treated with 20 µg/mL BNPs and incubated overnight before induction. Green: Dll4 mRNA expression. Scale bar: 100 µm. Quantification and comparison of cytoplasmic **(B)** and nucleus **(C)** Dll4 mRNA expression in MSCs under control and BNPs treated groups. **(D-E)** Cumulative probability distribution of cytoplasmic Dll4 expression of control and BNPs-treated groups, respectively. Error bars, s.e.m (n=4), with 100 −150 cells. p-Values were calculated using a two-sample t-test with respect to control. ns, not significant, *** p < 0.001.

Between the different days measured, there is no significant difference. For the BNPs-treated group, cytoplasmic Dll4 is significantly higher, particularly on day 3 and day 5, indicating a strong effect of BNPs. The data suggest that BNPs progressively enhance the cytoplasmic Dll4, with the most substantial effects observed by day 3. **Fig. 5C** showed the comparison of nucleus Dll4 expression over different days with different treatments. For the control group, Dll4 expression across day 1, 2, 3, and 5 showed relatively same levels with no statistically significant changes. In contrast, the BNP-treated group displays a gradual increase, with a significant rise by day 5, indicating the delayed but potent effect of BNPs. These data highlights BNPs’ time-dependent efficacy in enhancing osteogenic activity. Furthermore, for the cytoplasmic Dll4, we further conducted cumulative analysis to understand the trend and patterns of Dll4 expression over time, which could potentially reveal insights into the progression and impact of our interventions. **Fig. 5D-5E** showed the cumulative probability of Dll4 expression over several days (1, 2, 3, and 5 days) for control and BNPs-treated groups, respectively.

These results confirmed the effects of BNPs on Dll4 expression is time dependent. These data further support the importance of Notch signaling in regulating BNPs enhanced Dll4 expression during osteogenic differentiation, which demonstrates BNPs’ potent and sustained impact, important for fields like drug delivery or gene therapy.

## Discussion

In this study, we first investigated the effects of intracellularly uptake BNPs on osteogenic differentiation and examined the regulatory roles of Notch signaling in enhancing this differentiation process, utilizing an LNA/DNA nanobiosensor. This nanobiosensor, unlike conventional approaches for mRNA detection, allows for the monitoring of mRNA expression in live cells at the single-cell level without requiring cell lysis or fixation. This capability permits continuous observation of Dll4 mRNA gene expression dynamics throughout the process of osteogenic differentiation. The specificity and stability of this nanobiosensor has been demonstrated earlier ^4^. Previous studies have demonstrated that this nanobiosensor can track spatiotemporal RNA dynamics in collective cell migration ^39^, mice lung cancer ^40^, wounded corneal tissue repair ^41^, liver tissue ^42^, and vasculature formation ^35, 43^. Our group recently showed this nanobiosensor in monitoring Dll4 dynamics during osteogenic differentiation ^2, 4^. It is noted that this nanobiosensor can be utilized to detect other types of RNA detection, i.e., miRNA and long non-coding RNA (lncRNA) ^32, 34, 44, 45^. Moreover, this nanobiosensor is versatile, working effectively across various types and tissue environment. Previous studies have shown this nanobiosensor can monitor miRNA dynamics during 3D cancer invasion, mRNA and lncRNA dynamics during 3D osteogenic differentiation ^34, 46^. The ability to monitor gene expression in 3D physiological environments opens up possibilities to discover new aspects and mechanisms of cell-cell and cell-matrix interactions, ultimately paving the way for the development of innovative tools in tissue engineering and regenerative medicine.

Bone based ECM, derived from decellularized bone tissues, can be used as powder, hydrogel, and electrospun scaffolds in regenerative therapies for bone repairs and wound healing ^47–50^. These scaffolds exhibit robust mechanical properties, inherent osteoinductive and osteoconductive capabilities, and closely mimic natural bone. Although various studies have shown decellularized bone matrix promote bone regeneration, the effects of BNPs on tissue regeneration and its fundamental mechanism remains largely unknown. We recently reported these BNPs enhance bone repair *in vivo* ^24^. Due to their native source, BNPs exhibit unique characteristics, including biocompatibility, bioactivity, osteoinductivity, osteoconductivity, and biodegradability.

Moreover, these nanoparticles can be engineered to carry drugs, growth factors, or other therapeutic agents directly to the site of bone damage. Targeted drug delivery system can improve the effectiveness of treatments while reducing side effects associated with systemic drug delivery. Importantly, the properties of BNPs can be tailored during their synthesis process to meet specific requirements of different applications, such as varying their size, surface charge, and functionalization with bioactive molecules. These unique features of BNPs harness the natural properties of bone to offer promising solutions in bone tissue engineering, making them a focus of current research in regenerative medicine and related fields.

One of the primary reasons for the unique features of BNPs is the protein content they contain. We have shown these BNPs contain several major ECM proteins, including TGF-β, fibronectin, and COL1A1 ^24^. It is noted that the enhancement of cell proliferation and osteogenic differentiation can be observed quickly after several days of incubation, **Fig. 2** and **Fig. 3**. One reason is that BNPs can gradually release protein over 5-7 days due to the physical connections within the nanoparticles, without the need for crosslinking. The release profile has been studied previously ^24^. BNPs enhance cell proliferation and osteogenic differentiation may be related to the protein contained in the nanoparticles (i. e., TGF-β). Previous studies have shown that TGF-β promotes osteoinduction of osteoblasts and bone marrow stromal cells ^51–53^. The osteogenic differentiation process is modulated by several growth factors including bone morphogenetic proteins (BMPs), TGF-β, fibroblast growth factor (FGF), and platelet derived growth factor (PDGF). Several signal transduction pathways have been identified during osteogenic differentiation, including BMP signaling, Wnt/b-catenin signaling, and Notch signaling ^51–56^. Our previous studies showed Notch signaling is activated and required for osteogenic differentiation, inhibition of Notch signaling repress osteogenesis, with reduced ALP activity ^2, 4^.

Notch signaling is a highly conserved evolutionary pathway that influences cell proliferation, cell fate determination, and stem cell differentiation in both embryonic and adult tissues ^57–60^. This pathway includes four Notch receptors (Notch1-4) and five distinct ligands (Dll1, Dll3, Dll4, Jag1, and Jag2). Recently, the involvement of Notch signaling in osteogenic differentiation has gained significant attention from researchers, with various studies confirming its activation during this process ^36, 61^. Recently, it is reported that the Notch ligand Dll4 can promote bone formation in male mice without causing adverse effects in other organs ^62^. Another study reported that Notch signaling is crucial for regulating the differentiation and function of osteoblasts and osteoclasts, as well as maintaining skeletal homeostasis ^63^. Cao *et al*. demonstrated that the Notch receptor Notch1 and Notch ligand Dll1 play roles in osteogenic differentiation ^64^. Their study showed that inhibiting Notch1 decreased ALP activity during BMP-induced osteogenic differentiation of MSCs *in vitro*. However, the exact involvement of Notch signaling during BNPs enhanced osteogenic differentiation remains largely unexplored, especially the time-dependent of Notch involvement. In this study, we aim to characterize the Notch signaling pathway during spontaneous BNPs enhanced osteogenic differentiation during whole period of cell differentiation. To achieve this aim, we monitored and analyzed Notch ligand, Dll4 mRNA expression across different time points. Our results showed that Notch signaling is involved in BNPs enhanced osteogenic differentiation. Intracellular uptake BNPs in MSCs promote both early (1-7 days) and late stage (8-21 days) differentiation, as indicated by the increase in the production of both early (ALP activity) and late osteogenic markers (calcium mineralization). Interestingly, the analysis of the expression dynamics of Notch ligand, Dll4 expression has shown significant increase of cytoplasmatic Dll4 in BNPs treated MSCs during osteogenic differentiation process, suggesting intracellular uptake of BNPs could modulate gene expression profile. To further verify the involvement of Notch signaling, we investigated the effects of γ-secretase inhibitor DAPT on osteogenic differentiation. Without BNPs, disruption of Notch pathway decreased Dll4 expression, reduced ALP activity and calcium mineralization. The MSCs were treated with BNPs, the effects of Notch inhibition on osteogenic differentiation were mediated with enhanced Dll4 expression and increased calcium mineralization. Overall, our study suggests that Notch signaling in involved and regulate osteogenic differentiation. Internalization of BNPs enhanced osteogenic differentiation with increased Dll4 expression, indicating BNPs may activate Notch signaling. To our knowledge, only a few studies have investigated the Dll4 ligand during osteogenic differentiation and none in our cellular model in live cells ^55^. Moreover, Dll4 has been reported to be upregulated during endochondral and intramembranous bone regeneration ^54^.

Although it is mentioned in literature TGF-β promotes proliferation, early differentiation, and commitment to the osteoblastic lineage through the selective BMP, Wnt, Smad2/3, and Notch signaling ^55, 56^, it is unclear whether the enhancement of osteogenic differentiation is due to single or multiple protein molecules presented in the BNPs. Thus, further mechanistic studies are required to elucidate the molecular and cellular processes that regulate osteogenic differentiation. Specifically, the fundamental regulatory mechanisms of Notch pathway and its upstream and downstream signaling pathways should be further investigated using loss- and gain-of function experiments. Understanding the fundamental mechanisms of the osteogenic differentiation of MSCs induced by intracellular uptake of BNPs will provide valuable information that can be used for bone regeneration and repair.

### Conclusions

In this study, we investigated the effects of intracellular uptake of BNPs on cell proliferation and differentiation and the involvement of Notch signaling during osteogenic differentiation process. We first showed BNPs enhance MSCs proliferation and osteogenic differentiation, with significantly increased number of cells, ALP enzyme activity, and calcium deposition. By leveraging an LNA/DNA nanobiosensor, we examined and compared the Notch ligand, Dll4 mRNA expression dynamics in cytoplasm and nuclear during osteogenic differentiation process. Pharmacological disruption of Notch signaling using γ-secretase inhibitor DAPT mediated osteogenic process, with reduced expression of early and late stage of differentiation markers (ALP, calcium mineralization). The decrease in the expression of osteogenic markers in cells treated with DAPT suggests that Notch pathway is involved in their regulation. In addition, the expression of Dll4 in BNPs treated cells displayed a time-dependent profile, which is consistent with the enhancement of cell proliferation and differentiation. Our results suggest that the changes in BNPs treated cells during osteogenic differentiation may be linked to the increased expression in the Dll4 mRNA. In conclusion, this study will add new insights concerning the osteogenic differentiation of MSCs as well as the molecular mechanisms by which BNPs can stimulate the differentiation process. Results showed that BNPs induced effects on osteogenesis can be associated to the modulation of Notch signaling which plays essential role in cell fate and differentiation. Further studies will focus on elucidating the relationship among TGF-β, Notch, and BMP signaling pathways during osteogenic differentiation process as well as the effects of BNPs.

## Acknowledgment

S. Wang acknowledge the financial support from NSF CAREER (CMMI: 2143151).

## Author contributions

B. W. and S.W conceived the initial idea of the study. A. S., J. C. and S. F. performed the experiments. B. W. and S.W. contributed to the experimental design and data analysis. B. W. and S.W. wrote the manuscript with feedback from all authors.

## Data Availability Statement

The original contributions presented in the study are included in the article/Supplementary Material, further inquiries can be directed to the corresponding authors.

