## Supplementary material for "Notch Signaling in regulating Bone-derived Nanoparticles (BNPs) enhanced Osteogenic Differentiation": Fig. S1-S4

*Corresponding Author:

**Fig. S1.** Effects of internalized BNPs on cell viability.

**Fig. S2.** Comparison of ALP intensity of MSCs after 7 days of osteogenic induction

**Fig. S3** Images of MSCs in 12-well plate after 14 days of osteogenic induction

**Tab. S1.** LNA/DNA probes and quencher sequences

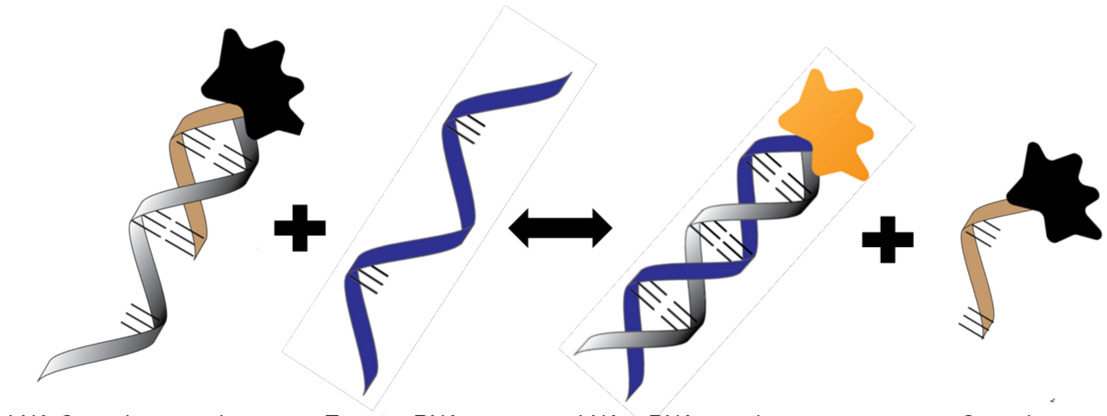


**Fig. S1 Illustration of LNA/DNA nanobiosensor**. The LNA/DNA nanobiosensor is a complex of LNA donor and quencher probe. The fluorophore at the 5' of LNA donor probe is quenched due to close proximity. In the presence of target mRNA sequence, the LNA donor sequence is displaced from the quencher to bind to the target sequence, allowing the fluorophore to fluorescence.

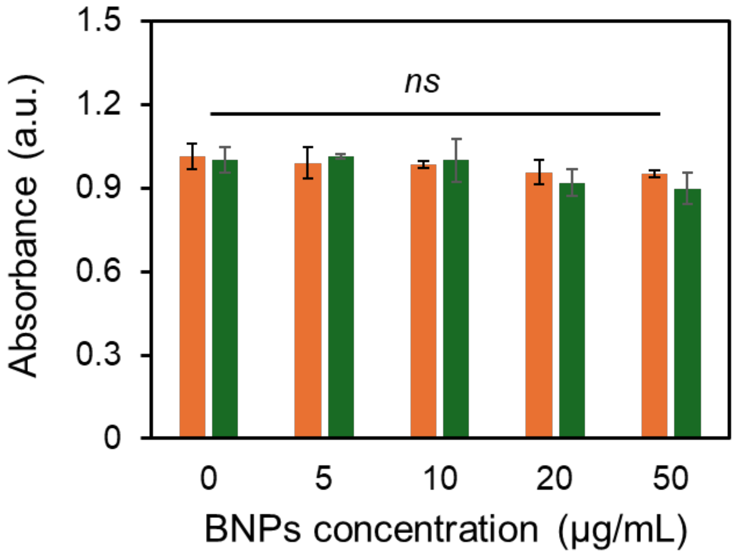


**Fig. S2 Effects of internalized BNPs on cell viability.** MSCs were treated with BNPs with the concentration of 0, 5, 10, 20, and 50 μg/mL. After 14 days of incubation, the absorbance at 450 nm was read using fluorescent microplate reader (BioTek, Synergy 2). Data expressed as mean± s.e.m. (n=3).

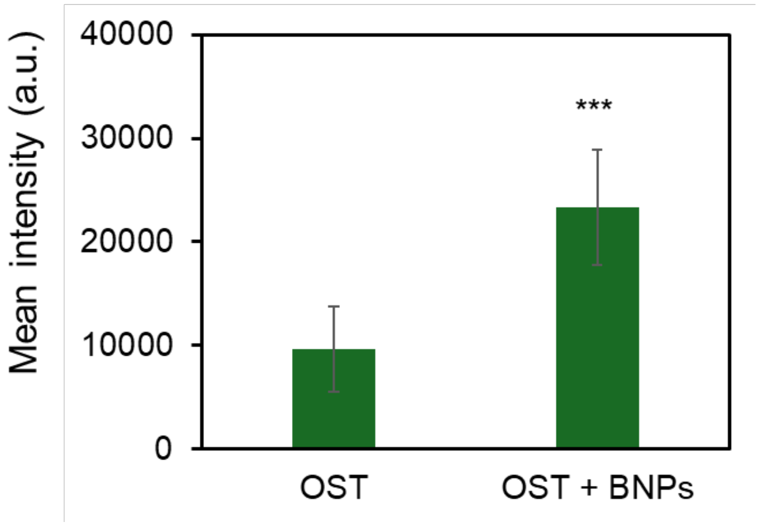


**Fig. S3** Comparison of ALP intensity of MSCs after 7 days of osteogenic induction. Cells were treated with BNPs (20 μg/mL) and cultured in osteogenic induction medium. At least 100 cells were analyzed for each condition. Data expressed as mean± s.e.m. (n=3).

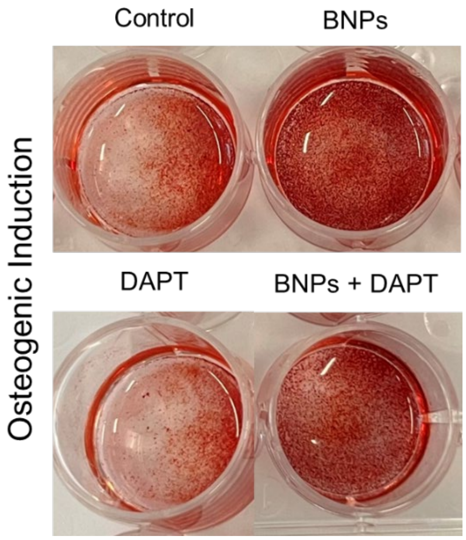


**Fig. S4** Images of MSCs in 12-well plate after 14 days of osteogenic induction. For BNPs treated groups, MSCs were treated with BNPs at the concentration of 20 μg/mL overnight for internalization. DAPT (20 μM) were added to MSCs for Notch inhibition.

**Tab. S1.** LNA/DNA probes and quencher sequences

| Name | | Sequence (5'-3') | Fluorophore |
|---|---|---|---|
| Dll4 mRNA | Donor | +AA +GG +GC +AG +TT +GG +AG +AG +GG +TT | /56-FAM |
| | Quencher | +TT +CC +CG +TC +AA | /3-Iowa BlackFQ |
| | Target | AA CC CT CT CC AA CT GC CC TT | |

* + represents LNA monomer
